# The Role of Macronutrient Composition in Semipurified Diets in Shaping Distinct Obesity Phenotypes: Evidence for Hepatic Oxidative Stress Involvement

**DOI:** 10.1101/2025.07.17.665389

**Authors:** Pedro Rocha Tenorio, Gabriel Smolak Sobieski e Silva, Isadora Chagas Vercellone, Débora Hipólito Quadreli, Juliany Carolina Duma de Castro, Glaura Scantamburlo Alves Fernandes, Fabio Goulart de Andrade

**Affiliations:** Department of Physiological Sciences, State University of Londrina, Londrina, Brazil; Department of Pathology Sciences, State University of Londrina, Londrina, Brazil; Department of General Biology, State University of Londrina, Londrina, Brazil; Department of Histology, State University of Londrina, Londrina, Brazil

**Keywords:** High-Fat Diet, High-Sugar Diet, Metabolic Syndrome, Diabetes, Metabolic Dysfunction-Associated Steatotic Liver Disease (MALSD)

## Abstract

While obesity is widely recognized as a global epidemic, the specific roles of food processing and macronutrient composition in driving metabolic dysfunction remain complex. Current animal models often rely on purified diets that may lack translational relevance to human dietary patterns. This study aimed to investigate how the macronutrient imbalances in purified diets influence obesity, metabolic comorbidities, and oxidative stress in adult rats. We compared a commercial grain base diet (C/GB) with two semi-purified diets (SD): a balanced SPD (B/SP) and a high-fat/high-sugar SPD (HFS/SP). The study evaluated body weight trajectories, adiposity distribution, glucose/lipid metabolism, and systemic and tissue-specific oxidative stress in male Wistar rats over a 10-week period. The HFS/SP group exhibited a unique biphasic weight trajectory, characterized by an initial rapid increase in body weight and hyperphagia during the first three weeks, followed by loss and stabilization. This was accompanied by significant dyslipidemia, impaired glucose regulation, hepatic steatosis, and elevated systemic oxidative stress. In contrast, the B/SP group showed continuous weight gain and increased adiposity with redistribution of adipose tissue toward subcutaneous depots, preserved glucose handling, and enhanced fat mobilization. Notably, significantly elevated reactive oxygen species production was localized primarily in the liver of the HFS/SP group. Our findings suggest that the nutritional imbalance in processed foods acts as a critical driver of metabolic disease. Rather than being driven solely by adiposity or calories, metabolic dysfunction is heavily influenced by dietary quality, where high-processing and imbalanced macronutrient intake trigger a rapid transition from compensatory weight gain to pathologically dysregulated metabolic syndrome, which is closely associated with hepatic oxidative stress.

## INTRODUCTION

The global prevalence of obesity has steadily increased since 1975, contributing to 8.9% of global mortality. Recognized as an epidemic by the World Health Organization in 1995, obesity is projected to affect 51% of the global population by 2030, posing one of the 21st century’s most pressing public health challenges^1^. The most widely accepted pathophysiological mechanism of obesity involves excessive adipose tissue accumulation coupled with impaired energy substrate oxidative capacity. This imbalance elevates reactive oxygen species production, primarily within mitochondria and cellular membranes. While initially localized, oxidative stress can become systemic via bloodstream dissemination. In insulin-sensitive organs like the liver, systemic oxidative stress disrupts metabolic homeostasis, impairs insulin signaling and promotes hypercholesterolemia. These disturbances establish a positive feedback loop driving steatotic liver disease and metabolic syndrome, including type 2 diabetes and cardiovascular disease^2–4^.

Animal models of diet-induced obesity (DIO) frequently employ purified diets composed of isolated proteins, refined carbohydrates, and purified fats to ensure high reproducibility. These diets often feature high caloric density from fat (e.g., 60% kcal) to facilitate rapid metabolic syndrome via lipotoxicity and oxidative stress^5^. However, such models may have limited translational relevance because they reduce the diet to a simplified macronutrient composition, without accounting for components of the food matrix, such as phytochemicals, that may modulate digestion and metabolism^5^. Furthermore, their extreme composition deviates from typical human dietary patterns^6^, yielding distinct metabolic profiles^7,8^. In humans, observational studies link ultra-processed food consumption to obesity and metabolic syndrome^9^, yet randomized intervention trials remain scarce and brief^10,11^. Similarly, rodent studies using purified diets yield divergent results: some indicate metabolic disturbances^12,13^, while others show no detrimental effects^14,15^ or even improvements over standard chow^16^. Semi-purified (SP) diets, combining natural and refined ingredients, offer greater translational relevance by closely reflecting human patterns^17^. Despite their translational potential, SP diet models remain underutilized. Crucially, there is a lack of evidence regarding how food processing and the resulting changes in the food matrix influence obesity and oxidative stress when macronutrient ratios mimic human consumption. To address this gap, we investigated the impact of food processing on obesity, comorbidities, and oxidative stress in adult rats by comparing a natural-ingredient diet with two SP diets: a balanced formulation and a high-fat/high-sugar formulation designed to mimic human dietary patterns.

## MATERIAL AND METHODS

### Animal Housing and Management

The primary outcome variable of this study was body weight gain. Given the lack of consensus on rodent obesity classification and observed inter-study variability in Wistar rat weight gain over comparable experimental periods, we adopted a conservative benchmark: an estimated 25% increase over the expected body weight gain of approximately 80±10 g was considered the minimum biologically relevant effect to be detected^17^. This corresponds to a mean additional weight gain of 20 g in adult animals, based on the characteristics of the animals housed in our vivarium. With α = 0.05 and statistical power set at 0.80, the minimum required sample size was calculated as n ≈ 4 animals per group^18^; therefore, five animals were included in each group Fifteen 90-day-old male Wistar rats were randomly assigned to three dietary groups (n = 5/group) using random sequence generation (RStudio): a control commercial grain-based diet (C/GB) (Nuvilab CR1®, Quimtia, Brazil), a balanced semi-purified diet (B/SP), and a high-fat/high-sugar semi-purified diet (HFS/SP). The animals were housed under control conditions (22 ± 2ºC, 12 h light/dark cycle) with *ad libitum* access to water and food for 73 days.

On the 74th day, following a 6-8 hour fast, between 02:00 and 04:00 pm, the animals were anesthetized via intraperitoneal administration of ketamine (100 mg/kg) and xylazine (10 mg/kg). Blood samples were collected from the inferior vena cava into tubes containing a coagulation activator (FristLab, Brazil) and the serum was stored in 20 °C. The liver, inguinal and retroperitoneal adipose tissues were dissected, washed in ice-cold PBS, fractioned, and fixed in 4% paraformaldehyde or snap frozen and stored at -80ºC.

### Diets Composition

The semi-purified diets were prepared weekly in the laboratory. Both semi-purified diets contained: 100 g soy protein isolate, 25 g wheat gluten, 25 g dried whole egg, 100 g maltodextrin, 100 g degermed cornmeal, 50 g wheat bran, 25 g psyllium husk, 100 g defatted coconut flour, 5 g Brazilian nut flour, 25 g brewer’s yeast, and 40 g of a mineral and vitamin mix. The B/SP diet was supplemented with 474 g of cornstarch, whereas the HFS/SP diet included 100 g of sucrose as added sugar and 155 g of lard. All ingredients were thoroughly mixed, oven-dried and stored in sealed containers. Nutrient composition was formulated according to NRC-95G recommendations to prevent micronutrient deficiencies. Because the HFS/SP diet had a higher energy density, fiber and micronutrient contents were adjusted on an energy basis to maintain comparable nutrient availability across diets and to minimize potential confounding effects of micronutrient inadequacy^19^. Detailed nutrient composition is presented in Table 1.

**Table 1.** Nutritional composition of diets.

| Nutritional Facts | C/GB | B/SP | HFS/SP |
| --- | --- | --- | --- |
| Crude Energy (kcal/kg) | 3468 | 3697 | 4625 |
| Carbohydrates (g/1000 kcal) (% kcal) | 150.81 (66) | 161.48 (71) | 78.70 (36) |
| Added Sugars (g/1000 kcal) (% kcal) | 0 (0) | 0 (0) | 25.30 (10) |
| Lipids (g/1000 kcal) (% kcal) | 11.53 (10) | 15.42 (14) | 54.92 (49) |
| Protein (g/1000 kcal) (% kcal) | 64.3 (24) | 40.84 (15) | 40.65 (15) |
| Fiber (g/1000 kcal) | 19.32 | 19.47 | 18.60 |
| Cholesterol (mg/1000 kcal) | 0 | 107.38 | 145.51 |
| Vitamin E (μg/1000 kcal) | 7.8 | 12.71 | 12.97 |

<sup>1</sup> **Legend:** C/GB = Commercial grain base diet; B/SP = Balanced semi-purified diet; HFS/SP = High-fat/high-sugar semi-purified diet.

### Murinometric and Dietary Parameters

Body weight, food intake, and water consumption were monitored weekly. Food and water intake were determined using the residual intake method and calculated at the cage level, normalized to the number of animals per cage. Weekly body weight and caloric intake data were used to calculate food efficiency (grams of body weight gain per kcal consumed per week)^12^. The adiposity index was calculated as the percentage of adipose tissue weight by body weight, while the visceral/subcutaneous adipose tissue ratio was calculated as the ratio of retroperitoneal to inguinal adipose tissue.

### Serum Metabolic Profile and Liver Markers

To assess glucose homeostasis, fasting glucose, the triglyceride-glucose (TyG) index, and fructosamine were measured. The TyG index was used as a surrogate marker associated with impaired glucose handling and metabolic syndrome^20^, whereas fructosamine was used as an indicator of medium-term glycemic control. Free fatty acids and glycerol were determined as indicators of fat mobilization, ketone bodies as markers of fat oxidation, and triglycerides and cholesterol as markers of lipid homeostasis. Corticosterone was quantified to account for its role in glucose and lipid regulation. Alanine aminotransferase (ALT), aspartate aminotransferase (AST), and gamma-glutamyl transferase (GGT) were assessed as indicators of liver function.

The TyG index was calculated as *ln* [**fasting triglycerides** *(mg/dl)*×**fasting glucose** *(mg/dl)* /2 ]). Fasting glucose, serum fructosamine, triglycerides, glycerol, total cholesterol, HDLc, albumin, ALT, AST, and GGT were measured using commercial kits (Vida Biotecnologia, Brazil). Free fatty acids were measured using diphenyl carbazide method^21^. Ketone body concentration was determined after serum deproteinization by the conversion of β-Hydroxy butyric acid into acetoacetic acid using John’s reagent and subsequent measurement with sodium nitro prussiate in an alkaline glycine solution^22^. Corticosterone concentrations were measured using NBT colorimetric method^23^.

### Tissues Histology, Metabolism, and Composition

Liver and adipose tissue samples were fixed, embedded in paraffin, sectioned at 5 μm, and stained with hematoxylin and eosin. Histological images were captured using a BA310MET-T light microscope (Motic, China) equipped with a Moticam A5 digital camera.

Twenty liver images from each animal, acquired at 10× magnification, were independently evaluated by two blinded experts. When a consensus was not achieved, a third evaluator reviewed the images to determine the final score. Histopathological diagnosis was performed according to the NASH Clinical Research Network Scoring System, classifying hepatic steatosis, hepatocellular ballooning, and inflammation based on established criteria^24^. Adipocyte diameter was calculated from a minimum of 500 adipocytes per sample using ImageJ software (v1.53, NIH) with the Adiposoft plugin (v1.16), and tissue cellularity was subsequently estimated^25^.

Total lipoprotein lipase (LPL) and heparin-released LPL activity, considered the functional fraction of the enzyme involved in lipid uptake, together with lipolysis capacity, were measured as previously described^26^ with minimal modifications. For heparin-released LPL, an additional incubation step with 5 U/mL heparin was included. Lipolysis was assessed using 2,3-dimercapto-1-propanol tributyrate as the substrate and 5,5-dithiobis-2-nitrobenzoic acid as the color reagent, 1 U = 1 *µ*M of free fatty acid released per minute.

To assess tissue composition, triglycerides, cholesterol, glycogen, and glucose were extracted from frozen liver samples using the Folch method, followed by digestion of residual tissue debris in 20% (w/v) KOH with heating^27^. Triglycerides, cholesterol, and glucose were quantified using commercial kits, whereas glycogen content was determined by the phenol–sulfuric acid method^28^.

### Oxidative Markers

Serum and homogenized tissue samples in PBS pH 7.4 were used to determine antioxidant capacity by ferric-reducing antioxidant power^**29**^ and total free thiols^**30**^. Oxidative stress markers included total reactive oxygen species (modified FOX-1 assay)^31^, lipid peroxidation^32^, and protein carbonylation^33^.

### Statistical Analysis

Residual normality and homoscedasticity were assessed using the Shapiro–Wilk and Brown–Forsythe tests, respectively. When both assumptions were satisfied, one-way ANOVA followed by Tukey’s HSD post hoc test was performed. In case of heteroscedasticity, Welch’s ANOVA followed by the Games–Howell post hoc test was used. When residuals deviated from normality, logarithmic transformation was applied. When the assumptions were subsequently satisfied, conventional or Welch’s ANOVA was applied, as appropriate. If normality was violated after transformation, the Kruskal–Wallis test followed by Dunn’s multiple-comparisons test was used.

Body weight was analyzed using the individual animal as the experimental unit, whereas food intake, water intake, caloric intake, and food efficiency were analyzed using the cage as the experimental unit, as these variables were measured at the cage level. When the assumption of sphericity was violated, the Greenhouse–Geisser correction was applied. Partial eta-squared was reported as an effect size for the two-way ANOVA, while Hedges’ g and corresponding 95% confidence intervals were reported for pairwise comparisons. All statistical analyses were performed using RStudio (Posit, USA), with the onewaytests, effectsize, and ggplot2 packages. Statistical significance was set at p < 0.05.

## RESULTS

### Murinometric and Dietary Parameters

Initial body weights were comparable across groups. The B/SP group exhibited steady weight gain, resulting in the highest final body weight among all groups. In contrast, the HFS/SP group demonstrated a unique biphasic weight trajectory, characterized by an initial rapid increase in body weight followed by weight loss and subsequent stabilization (Fig. 1a). While weekly caloric intake did not differ significantly between groups, there were significant differences in the timing of consumption; both semi-purified diets showed higher initial caloric intake, followed by a normalization in the B/SP group and a non-significant reduction in the HFS/SP group (Fig. 1b). Food efficiency was significantly higher in the B/SP group compared to all other groups (Fig. 1c). Additionally, water consumption was reduced in both semi-purified diet groups (Fig. 1d).

**Figure 1.**
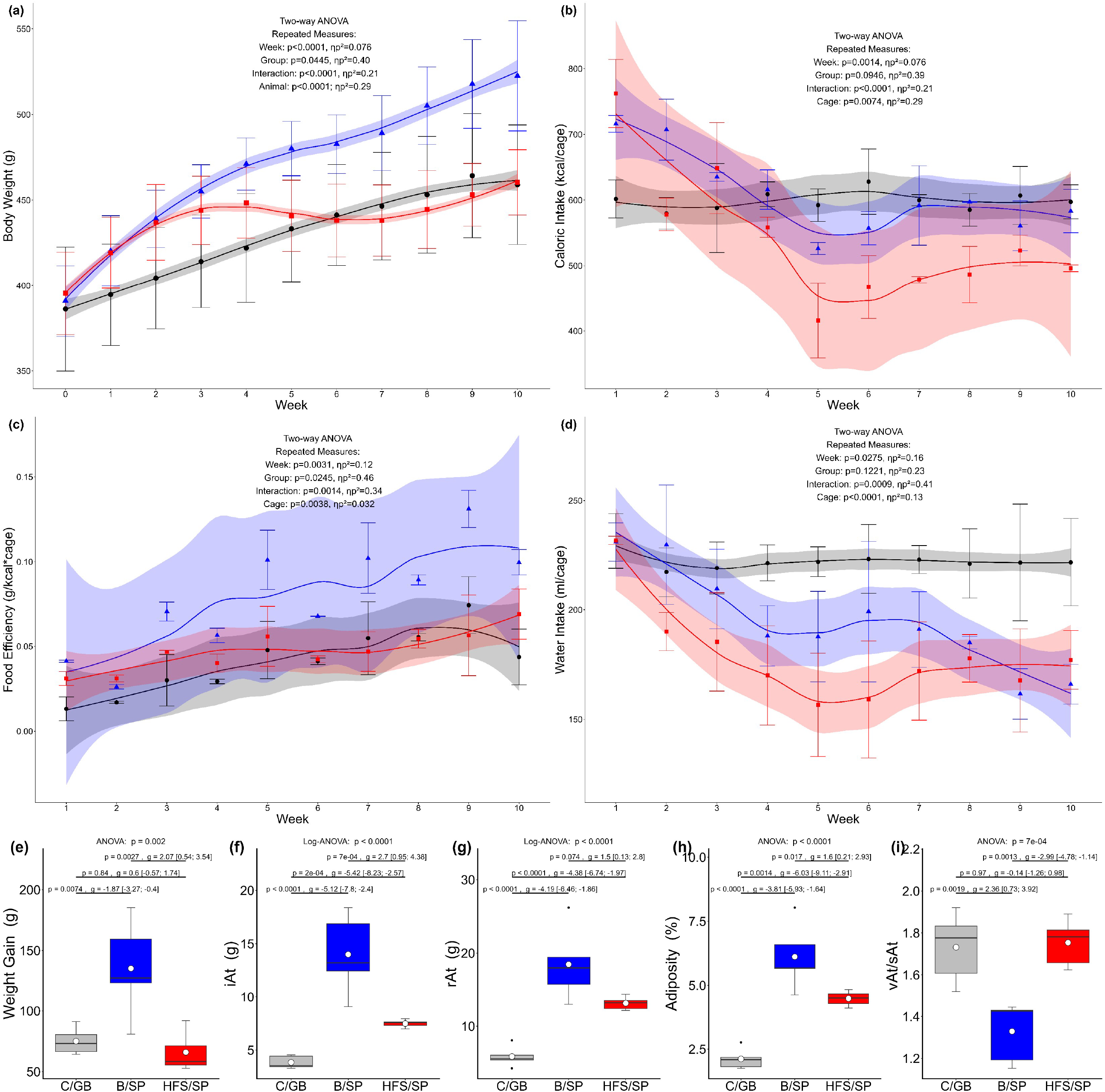
Effects of dietary interventions on dietary and morphometric parameters: Body weight gain throughout time (a), weekly caloric consumption per cage normalized by the number of animals (b), weekly food efficiency in g gain by kcal consumed per cage (c), weekly water consumption per cage normalized by the number of animals (d), body weight gain per animal (e), inguinal adipose tissue weight (f), retroperitoneal adipose tissue weight (g), adiposity index (h), and visceral to subcutaneous adiposity index (i), Diet groups: C/GB = commercial grain-based diet (⍰), B/SP = balanced semi-purified diet (▲), HFS/SP = high-fat high-sugar semi-purified diet (■). Data are presented as mean and standard deviation plus regression line and shadow area for 95% Confidence interval and box plots and interquartile ranges with white dots indicating group means. Statistical analysis: Two-way ANOVA with repeated measures and partial eta-square effect size or One-way ANOVA with Tukey’s HSD post and Hedge’s g effect size with the 95% confidence interval.

Body weight gain was significantly higher in the B/SP group compared to both C/GB and HFS/SP, with no significant difference observed between the C/GB and HFS/SP groups (Fig. 1e). Marked differences in adipose tissue weight and distribution were observed across dietary interventions. The weights of subcutaneous and visceral adipose tissues, as well as the adiposity index, were significantly higher in animals fed semi-purified diets, with the greatest increase observed in the B/SP group (Fig. 1f-h). Furthermore, the visceral/subcutaneous adipose ratio was significantly lower in the B/SP group compared to the other two groups, indicating a shift toward subcutaneous fat deposition (Fig. 1i).

### Serum Metabolic Profile and Liver Markers

Markers of glucose homeostasis, including fasting glucose, fructosamine, and the TyG index, were significantly elevated in the HFS/SP group. Lipid metabolism was also profoundly altered in this group, characterized by significantly increased triglycerides, total cholesterol, and corticosterone levels, alongside a reduction in glycerol, free fatty acids, and ketone bodies. Conversely, the B/SP diet modulated lipid metabolism by decreasing triglyceride levels and increasing circulating glycerol, free fatty acids, ketone bodies, and total cholesterol. Regarding hepatic injury, markers were generally reduced in both semi-purified diets compared to the control; however, alanine aminotransferase (ALT) was significantly elevated in the HFS/SP group compared to the B/SP group (Table 2).

**Table 2.**
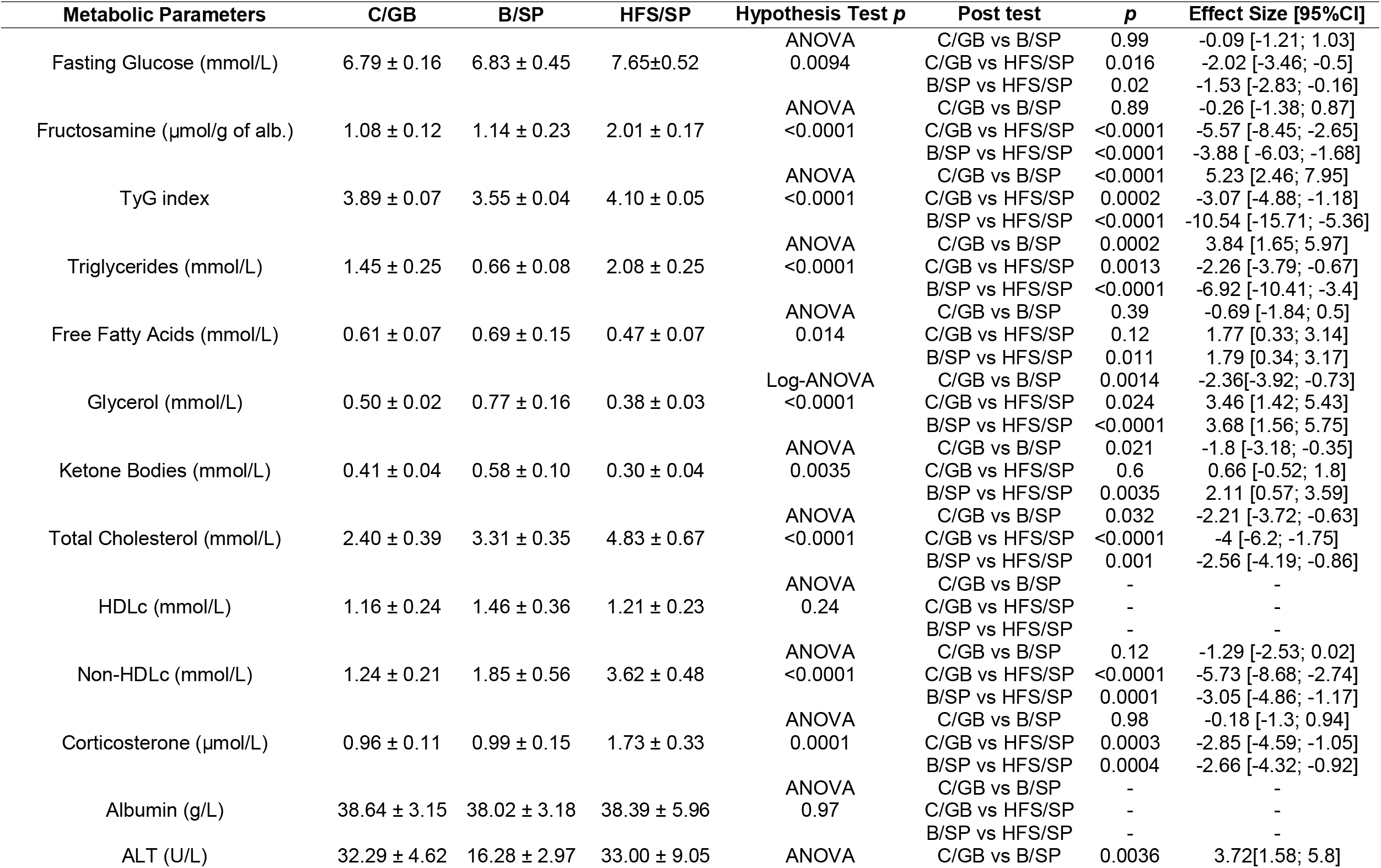
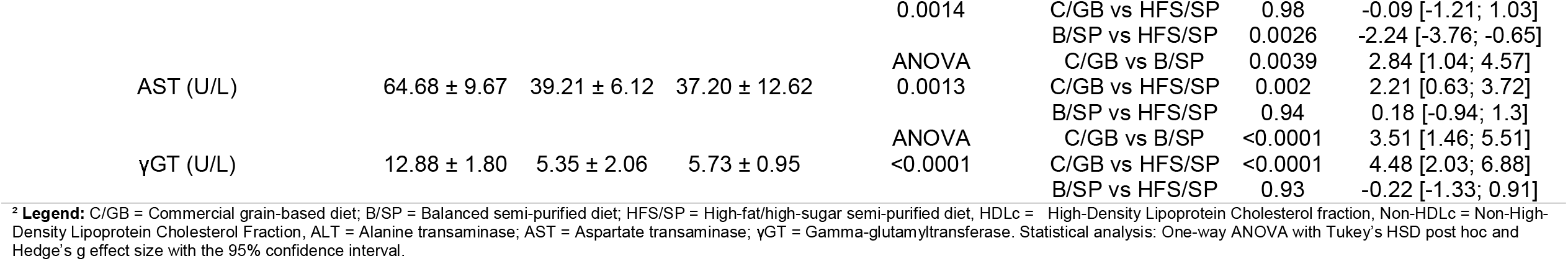
Serum metabolic panel.

### Tissues Histology, Metabolism and Composition

Semi-purified diets promoted adipose tissue accumulation at both sites; however, the relative weight of retroperitoneal and inguinal adipose tissue was significantly higher in the B/SP group compared to the HFS/SP group (Fig. 2a, g). Histological analysis revealed similar adipocyte hypertrophy in animals fed both semi-purified diets regardless of composition. In contrast, only the HFS/SP diet altered tissue cellularity, inducing adipocyte hyperplasia in retroperitoneal adipose tissue and a non-significant reduction in adipocyte number in inguinal adipose tissue (Fig. 2b, c, h, i). In retroperitoneal adipose tissue, both total LPL and heparin-released LPL fractions were elevated in HFS/SP animals, accompanied by reduced lipolysis, whereas the B/SP group exhibited increased lipolysis (Fig. 2d–f). In inguinal adipose tissue, the B/SP group showed elevated total LPL without a corresponding increase in heparin-released LPL or lipolysis; conversely, the HFS/SP group exhibited an increase only in heparin-released LPL, accompanied by reduced lipolysis (Fig. 2j–l).

**Figure 2.**
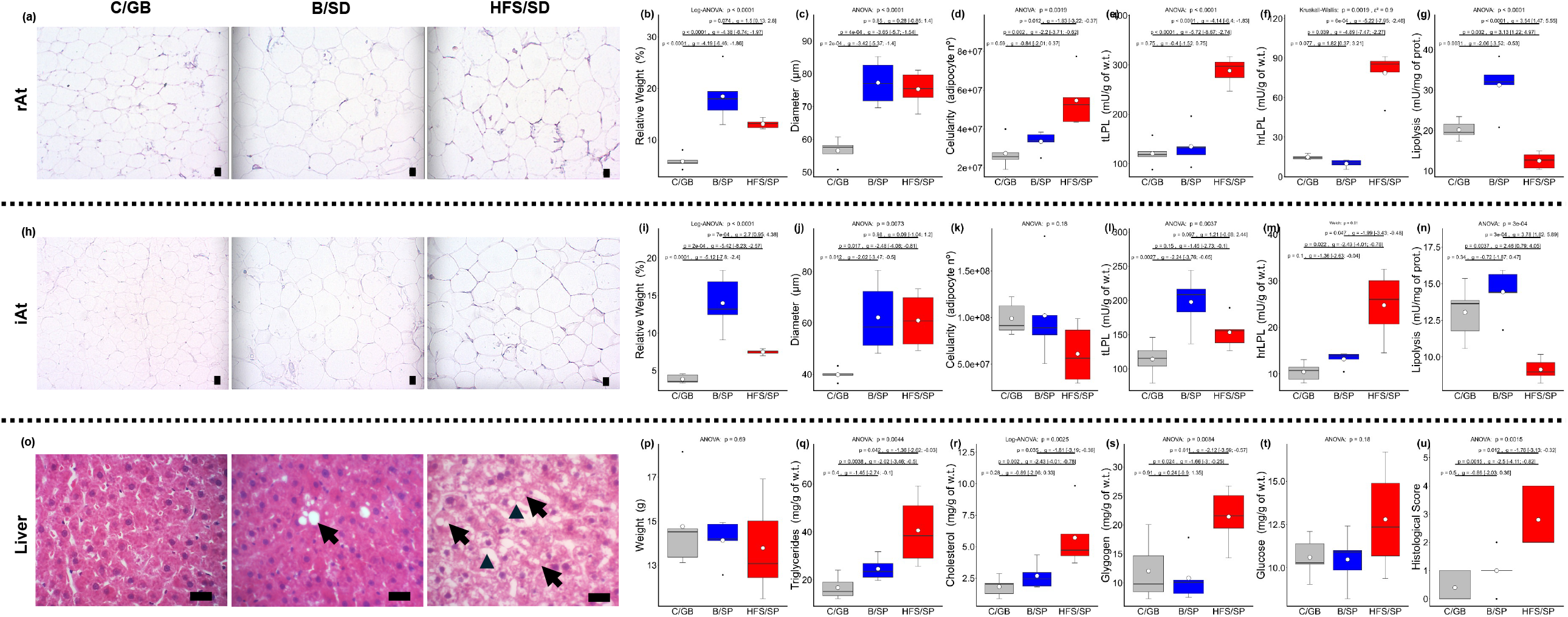
Histology, biochemical, and metabolic evaluation of adipose tissues and liver. Representative hematoxylin and eosin-stained retroperitoneal adipose tissues (a), inguinal adipose tissue (h), and liver sections (o); relative weight of retroperitoneal adipose tissue (b), inguinal adipose tissue (i), and liver (p); mean adipocyte diameter in retroperitoneal (c), and inguinal adipose tissue (j); estimated adipocyte number in retroperitoneal (d), and inguinal adipose tissue (k); total lipoprotein lipase activity in retroperitoneal (e), and inguinal adipose tissue (l); heparin-released fraction of lipoprotein lipase in retroperitoneal (f), and inguinal adipose tissue (m); in vitro lipolysis activity in retroperitoneal (g), and inguinal adipose tissue (n); hepatic triglyceride content (q); hepatic cholesterol content (r); hepatic glycogen content (s); hepatic glucose content (t); total histological lesion score of the liver(u). Diet groups: C/GB = commercial grain-based diet, B/SP = balanced semi-purified diet, HFS/SP = high-fat high-sugar semi-purified diet. Statistical analysis: One-way ANOVA with Tukey’s HSD post hoc, Welch’s ANOVA with Games–Howell correction or Kruskal-Wallis with Dunn’s multiple comparisons post hoc and Hedge’s g effect size with the 95% confidence interval. Data are presented as box plots as median and interquartile ranges with white dots indicate group means. Scale bar = 25 μm; (↖) arrows indicate lipid droplet, (▲) arrowhead indicates ballooning.

Liver weight did not differ among groups; however, biochemical analysis revealed a significant accumulation of triglycerides, cholesterol, and glycogen, along with a non-significant increase in glucose, specifically in the HFS/SP group (Fig. 2m–q). Histological analysis corroborated these findings, showing extensive mixed macrovesicular and microvesicular steatosis and hepatocyte ballooning in HFS/SP animals, resulting in a significantly higher histological score (Fig. 2r).

### Oxidative Markers

In the serum, semi-purified diets enhanced antioxidant capacity to a similar extent regardless of diet composition (Fig. 3a). However, oxidative markers were elevated specifically in the HFS/SP group compared to both C/GB and B/SP (Fig. 3b–e). In the liver, antioxidant capacity was reduced, and oxidative markers were significantly increased in animals fed the HFS/SP diet (Fig. 3f–j). In retroperitoneal tissue, while antioxidant capacity remained unaffected by diet, reactive species production was reduced in both semi-purified groups; however, oxidative stress markers were elevated specifically in the HFS/SP group (Fig. 3k–o). In inguinal adipose tissue, both antioxidant capacity and reactive species production were reduced in the semi-purified groups, whereas oxidative stress markers were significantly elevated in the HFS/SP group. Notably, oxidative status, assessed by total free thiols, was improved in the B/SP diet (Fig. 3p–t).

**Figure 3.**
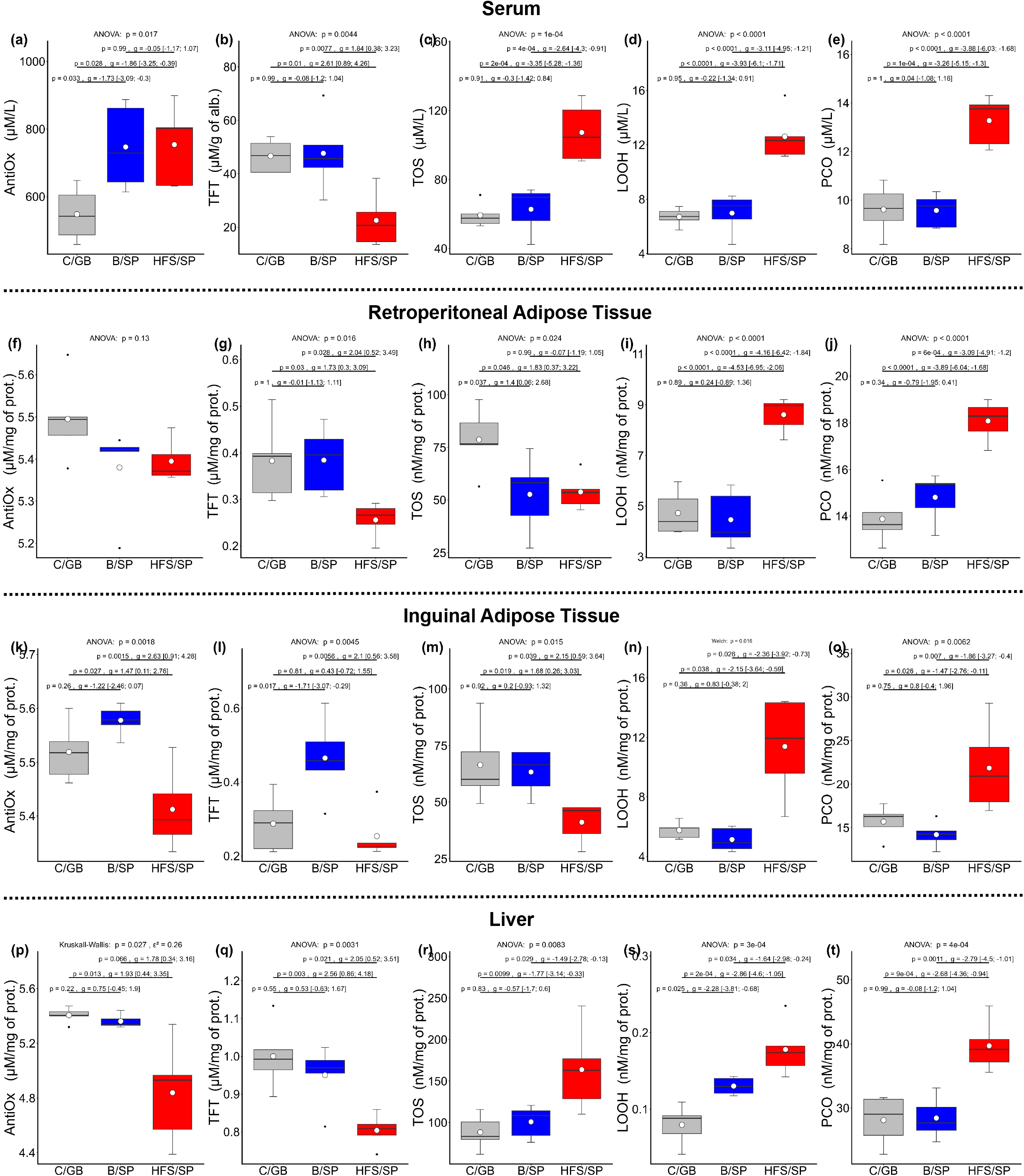
Oxidative markers in the serum, liver, and adipose tissues in animals fed grain-based and semi purified diets. Antioxidant capacity measured by ferric-reducing antioxidant power in ascorbic acid equivalent (AntiOx.) (a, f, k, and p), total free thiols (TFT) (b, g, l, and q), total oxidative status in equivalent of H2O2 (TOS) (c, h, m, and r), lipid peroxidation in equivalent of H2O2 (LOOH) (d, l, n, and s), and protein carbonylation (PCO) (e, j, o, and t). Diet groups: C/GB = commercial grain-based diet, B/SP = balanced semi-purified diet, HFS/SP = high-fat high-sugar semi-purified diet. Statistical analysis: One-way ANOVA with Tukey’s HSD post hoc, Welch’s ANOVA with Games–Howell correction or Kruskal-Wallis with Dunn’s multiple comparisons post hoc and Hedge’s g effect size with the 95% confidence interval. Data are presented as box plots as median and interquartile ranges with white dots indicate group means.

## DISCUSSION

The metabolic and phenotypic outcomes observed in this study demonstrate that dietary profiles can produce distinct metabolic trajectories beyond differences in caloric intake. The two semi-purified diets, which differed from the commercial grain-based diet in macronutrient composition, produced markedly different phenotypic, metabolic, and oxidative profiles, highlighting the importance of considering the combined nutritional characteristics of a diet when evaluating diet-induced metabolic dysfunction.

A notable feature of this study was the biphasic weight trajectory observed in the HFS/SP group. This was characterized by an initial period of hyperphagia and rapid weight gain during the first three weeks, followed by a transient loss of excess weight with reduction of food intake and a subsequent return to a normal weight-gain trajectory. Although food intake was evaluated at the cage level to avoid single-housing stress, potentially introducing cage-level cluster effects, the magnitude of the hyperphagic response was consistent across replicates. We believe that this initial spike reflects a failure to immediately compensate for the high energy density of the diet; while rodents are often described as hyperphagic when consuming high fat diets, they have been shown to attempt to adjust food intake in response to increased caloric density^7^. However, unlike standard DIO models, which fail to fully compensate for excess dietary energy, our results suggest that the HFS/SP group eventually engaged homeostatic mechanisms. This response may be related to the lower proportion of dietary fat in the HFS/SP diet compared with conventional DIO models^34^. Specifically, reduced fat mobilization and oxidation suggest partial preservation of visceral fat insulin signaling; however, tissue-specific signaling cannot be confirmed from the present data and warrants further investigation. We speculate that the decreased energy availability for skeletal muscle, combined with hypercortisolemia, resulted in skeletal muscle loss that prevented further weight gain^35^.

The divergence from traditional DIO models, which typically produce continuous, unchecked weight gain, likely stems from our use of a more human-like macronutrient profile. Indeed, previous work has shown that when dietary protein intake is high (≥15% kcal), only when fat intake accounts for 40% of kcal or greater do animals exhibit significant extra weight gain, even in obesity-prone lineages like C57BL/6 mice^36^. By utilizing lower fat concentrations than standard rodent DIO diets, we captured an initial weight spike that is consistent with the short-term weight gains reported in recent human intervention trials^10,11^. However, whereas those human trials are limited by their brief duration (1–2 weeks) and cannot capture long-term trends, our model allows for the observation of how these initial metabolic shifts may eventually trigger homeostatic stabilization.

In contrast, the B/SP diet promoted steady and significant weight gain. While carbohydrate-rich diets typically fail to induce obesity without additional metabolic challenges^37^, our data demonstrates that *ad libitum* intake of a semi-purified, carbohydrate-rich diet can indeed trigger obesity, possibly through increased metabolizable energy from the processed purified diet^38^. In the absence of an imbalanced composition, the metabolic alterations induced by diet purification appear to be compensatory. We observed increased fat mobilization from visceral adipose tissue and a redistribution toward subcutaneous depots, which may reduce the metabolic burden on internal organs and mitigate the metabolic dysfunction typically associated with greater visceral adiposity^39^.

Regarding specific nutrient content and its metabolic effects, the sugar-induced metabolic disturbances in glucose homeostasis are primarily associated with fructose liver spillover and their capacity to trigger hepatic metabolic dysregulation and steatosis. Our findings are consistent with this, as fructose was present exclusively in the HFS/SP diet through the addition of sucrose^3,40^. Dietary cholesterol is another component known for its implications in dysfunctional glucose handling via hepatic steatosis^41,42^; however, the levels used in this study (0.04% and 0.07%) were significantly below the concentrations typically required to induce metabolic dysfunction in rodents (0.5–1%). Furthermore, although the commercial diet differed from the experimental diets in its protein content, all groups provided sufficient protein to exceed NRC-95 recommendations for rat maintenance (>5% kcal), with all formulations being characterized as high-protein diets^19^. Previous work has shown that a 15% kcal protein intake minimizes energy intake in both low- and high-fat diets and exerts the maximal protective effect over obesity in high-fat diets without additional benefits from higher protein concentrations^36^. Consequently, we consider these nutritional discrepancies unlikely to be significant confounding factors in our metabolic observations.

A central mechanism associated with the observed comorbidities appears to be oxidative stress. Although systemic antioxidant capacity was increased in both semi-purified diet groups, possibly reflecting their higher vitamin E content and more standardized micronutrient composition relative to the grain-based diet, which has been shown to lack sufficient vitamin E to maximize antioxidant defenses^43^, this enhanced antioxidant capacity was insufficient to prevent oxidative damage in the HFS/SP group. We speculate that potent pro-oxidant stimuli, including fructose-induced mitochondrial ROS overproduction and lipid-induced cellular stress, may have overwhelmed these antioxidant defenses^3,44^. The elevated fructosamine levels observed in the HFS/SP group further support a potential contribution of non-enzymatic protein glycation and subsequent advanced glycation end-product formation to oxidative stress^45,46^. Notably, the liver was the only tissue exhibiting significantly increased reactive species production, indicating an association between hepatic oxidative stress and the systemic metabolic alterations observed in HFS/SP animals. This hepatic pattern may also help explain the relatively metabolically protected phenotype observed in the B/SP group. In the absence of significant macronutrient-induced hepatic steatosis and oxidative stress, these animals were able to gain weight without developing the systemic metabolic dysfunction observed in the HFS/SP group, consistent with human findings indicating that metabolic perturbations typically develop only once the liver is significantly affected^47,48^.

The partial dissociation between adiposity and metabolic complications observed in this study suggests that increased weight gain does not necessarily lead to equivalent metabolic deterioration, a distinction relevant to understanding human obesity subtypes. The B/SP group exhibited increased adiposity alongside a relatively preserved metabolic profile, resembling aspects of metabolically healthy obesity (MHO), in which excess fat mass is not necessarily accompanied by overt metabolic dysfunction and adipose tissue distribution may influence metabolic risk. This interpretation is consistent with observational evidence linking MHO in young adults to a more favorable systemic oxidative status compared with metabolically unhealthy obesity^49^. These findings raise the possibility that hepatic oxidative alterations may accompany the transition toward a metabolically unhealthy phenotype, although our results cannot establish whether hepatic oxidative stress contributes to metabolic deterioration or develops in parallel with it. Given evidence that MHO may be a transient state that progresses toward metabolic complications^39,50^, the MHO-like phenotype observed in our 10-week model may provide a useful framework for investigating this transition in future studies, although its temporal progression was not assessed here. Nevertheless, the contrast between the B/SP and HFS/SP groups suggests that the metabolic consequences of weight gain may depend not only on adiposity but also on dietary composition. The more pronounced hepatic oxidative alterations and metabolic disturbances in the HFS/SP group support an association between dietary imbalance, hepatic oxidative stress, and metabolic dysfunction, without establishing a causal relationship.

These findings suggest that the metabolic consequences of obesity may depend not only on adipose expansion but also on the development of hepatic alterations. While traditional models emphasize the progressive contribution of chronic adiposity to systemic oxidative stress and inflammation^2–4^, the HFS/SP group exhibited pronounced metabolic disturbances alongside hepatic steatosis and oxidative stress, despite relatively limited additional body-weight gain. Conversely, the B/SP group accumulated substantial adiposity without comparable hepatic or systemic metabolic alterations. This contrast is consistent with human evidence linking hepatic steatosis to the emergence of metabolic complications in obesity^35^, suggesting that hepatic involvement may help distinguish metabolically compensated adiposity from a metabolically unhealthy phenotype. Together, these findings position the liver as a potential link between dietary composition and the metabolic consequences of obesity, warranting further investigation into the contribution of hepatic oxidative stress to this relationship.

Regarding human relevance, or findings support considering dietary composition alongside energy intake and adiposity when evaluating metabolic health. The divergent phenotypes observed under the two semi-purified diets suggest that similar patterns of weight gain suggest that similar patterns of weight gain are not necessarily accompanied by equivalent metabolic outcomes. This interpretation is consistent with nutritional recommendations emphasizing balanced dietary patterns and minimally processed foods. Whoever, the present study does not isolate food processing from macronutrient composition or establish that either factor independently outweighs energy balance in preventing human obesity and metabolic disease. Rather, our findings provide an experimental basis for investigating how dietary composition may modify the metabolic consequences of excess adiposity.

## STUDY LIMITATIONS

Several limitations should be noted. First, because we did not employ a full factorial design, the independent effects of diet processing and macronutrient composition cannot be statistically isolated. Second, the sample size was powered specifically for body weight gain as the prespecified primary outcome; therefore, statistical power for secondary outcomes may have been limited. Third, the present findings do not establish a causal role for hepatic oxidative stress, and further mechanistic studies are required to determine whether modulation of hepatic oxidative pathways can prevent or reverse the associated metabolic alterations. Finally, using exclusively male rats, the influence of sexual dimorphism on nutritional responses remains to be investigated.

## CONCLUSION

In summary, the present study demonstrates that different dietary profiles produce distinct obesity phenotypes in adult Wistar rats. A balanced semi-purified diet promoted greater body-weight gain and adiposity while being associated with adaptive changes in lipid handling, preserved metabolic homeostasis, and limited oxidative damage. Conversely, a high-fat/high-sugar semi-purified diet induced dyslipidemia, impaired glucose regulation, hepatic steatosis, and systemic oxidative stress despite lower caloric intake and minimal weight gain. These findings indicate that metabolic dysfunction is not necessarily proportional to adiposity and may be strongly influenced by the overall dietary profile, including its macronutrient composition and purified characteristics. The liver serves as a prominent site of oxidative imbalance, supporting an association between hepatic oxidative stress and metabolic dysfunction in this phenotype.

## AUTHOR CONTRIBUTIONS

Tenorio, P.R. was responsible for conceptualization, data curation, formal analysis, funding acquisition, investigation, methodology, visualization and writing of original draft. Silva, G.S.S.; Quadreli, D.H.; Vercellone, I.C.; Castro, J.C.D., were responsible for investigation. Fernandes, G.S.A. was responsible for resources. Andrade, F.G. was responsible for project administration, resources, supervision, validation and review and editing.

## STATEMENTS AND DECLARATIONS

### Ethical considerations

The animal study protocol was conducted in strict adherence to the Ethical Principles in Animal Research endorsed by the Brazilian College of Animal Experimentation and was approved by the Ethics Committee on Animal Use of the State University of Londrina (Official Letter Nº 074/2022 Protocol Nº 032.2022)

### Data Availability

The datasets generated and/or analyzed during the present study are not publicly available, but they are available from the corresponding author on reasonable request.

## REFERENCES

1. Finkelstein EA, Khavjou OA, Thompson H, et al. Obesity and severe obesity forecasts through 2030. Am J Prev Med 2012; 42: 563–70.

2. Jin X, Qiu T, Li L, et al. Pathophysiology of obesity and its associated diseases. Acta Pharm Sin B 2023; 13: 2403–2424.

3. Ahmed B, Sultana R, Greene MW. Adipose tissue and insulin resistance in obese. Biomed Pharmacother 2021; 137: 111315.

4. Garvey WT. Is Obesity or Adiposity-Based Chronic Disease Curable: The Set Point Theory, the Environment, and Second-Generation Medications. Endocr Pract 2022; 28: 214–222.

5. Shahidi F, Pan Y. Influence of food matrix and food processing on the chemical interaction and bioaccessibility of dietary phytochemicals: A review. Crit Rev Food Sci Nutr 2022; 62: 6421–6445.

6. U.S. Department of Agriculture, Agricultural Research Service. Table 41. Nutrient Intakes per 1000 kcal from Food and Beverages: Mean Energy and Mean Nutrient Amounts per 1000 kcal Consumed per Individual, by Gender and Age, in the United States, 2017-2018. Beltsville (MD), 2020.

7. Bastías-Pérez M, Serra D, Herrero L. Dietary Options for Rodents in the Study of Obesity. Nutrients 2020; 12: 3234.

8. Speakman JR. Use of high-fat diets to study rodent obesity as a model of human obesity. Int J Obes (Lond) 2019; 43: 1491–1492.

9. Valicente VM, Peng C-H, Pacheco KN, et al. Ultraprocessed Foods and Obesity Risk: A Critical Review of Reported Mechanisms. Adv Nutr 2023; 14: 718–738.

10. Hamano S, Sawada M, Aihara M, et al. Ultra-processed foods cause weight gain and increased energy intake associated with reduced chewing frequency: A randomized, open-label, crossover study. Diabetes Obes Metab 2024; 26: 5431–5443.

11. Hall KD, Ayuketah A, Brychta R, et al. Ultra-Processed Diets Cause Excess Calorie Intake and Weight Gain: An Inpatient Randomized Controlled Trial of Ad Libitum Food Intake. Cell Metab 2019; 30: 67-77.e3.

12. Cheng HS, Phang SCW, Ton SH, et al. Purified ingredient-based high-fat diet is superior to chow-based equivalent in the induction of metabolic syndrome. J Food Biochem 2019; 43: e12717.

13. Takahashi E, Ono E. Effects of semi-purified diet on depressive behaviors in aged mice. Biochem Biophys Rep 2021; 28: 101152.

14. Almeida-Suhett CP, Scott JM, Graham A, et al. Control diet in a high-fat diet study in mice: Regular chow and purified low-fat diet have similar effects on phenotypic, metabolic, and behavioral outcomes. Nutr Neurosci 2019; 22: 19–28.

15. Del Bas JM, Caimari A, Ceresi E, et al. Differential effects of habitual chow-based and semi-purified diets on lipid metabolism in lactating rats and their offspring. Br J Nutr 2015; 113: 758–69.

16. Zhang L, Li X, Liu X, et al. Purified diet versus whole food diet and the inconsistent results in studies using animal models. Food Funct 2022; 13: 4286–4301.

17. Preguiça I, Alves A, Nunes S, et al. Diet-induced rodent models of obesity-related metabolic disorders-A guide to a translational perspective. Obes Rev 2020; 21: e13081.

18. Charan J, Biswas T. How to calculate sample size for different study designs in medical research? Indian J Psychol Med 2013; 35: 121–6.

19. National Research Council (US) Subcommittee on Laboratory Animal Nutrition. Nutrient Requirements of Laboratory Animals: Fourth Revised Edition, 1995. Washington, D.C.: National Academies Press (US). Epub ahead of print 1 January 1995. DOI: 10.17226/4758.

20. Wan H, Cao H, Ning P. Superiority of the triglyceride glucose index over the homeostasis model in predicting metabolic syndrome based on NHANES data analysis. Sci Rep 2024; 14: 15499.

21. Konobu K, Takao H. An improved colorimetric micromethod with diphenylcarbazide for serum free fatty acids determination. Yakugaku Zasshi 1978; 98: 226–30.

22. Essak Khan, Pranali Karanjkar, VN Ravi Kishore Vutukuri. Novel quantitative assay for estimation of ketone bodies in diabetic urine. Int J Sci Eng Res 2016; 7: 701–705.

23. Tu E, Pearlmutter P, Tiangco M, et al. Comparison of Colorimetric Analyses to Determine Cortisol in Human Sweat. ACS Omega 2020; 5: 8211–8218.

24. Ampuero J, Aller R, Gallego-Durán R, et al. The biochemical pattern defines MASLD phenotypes linked to distinct histology and prognosis. J Gastroenterol 2024; 59: 586–597.

25. Eriksson-Hogling D, Andersson DP, Bäckdahl J, et al. Adipose tissue morphology predicts improved insulin sensitivity following moderate or pronounced weight loss. Int J Obes (Lond) 2015; 39: 893–8.

26. Pérez-Torres I, Gutiérrez-Alvarez Y, Guarner-Lans V, et al. Intra-Abdominal Fat Adipocyte Hypertrophy through a Progressive Alteration of Lipolysis and Lipogenesis in Metabolic Syndrome Rats. Nutrients; 11. Epub ahead of print 5 July 2019. DOI: 10.3390/nu11071529.

27. Carr TP, Andresen CJ, Rudel LL. Enzymatic determination of triglyceride, free cholesterol, and total cholesterol in tissue lipid extracts. Clin Biochem 1993; 26: 39–42.

28. Schaubroeck KJ, Leitner BP, Perry RJ. An optimized method for tissue glycogen quantification. Physiol Rep 2022; 10: e15195.

29. Benzie IF, Strain JJ. The ferric reducing ability of plasma (FRAP) as a measure of ‘antioxidant power’: the FRAP assay. Anal Biochem 1996; 239: 70–6.

30. Koning AM, Meijers WC, Pasch A, et al. Serum free thiols in chronic heart failure. Pharmacol Res 2016; 111: 452–458.

31. Erel O. A new automated colorimetric method for measuring total oxidant status. Clin Biochem 2005; 38: 1103–11.

32. Jiang ZY, Hunt J V, Wolff SP. Ferrous ion oxidation in the presence of xylenol orange for detection of lipid hydroperoxide in low density lipoprotein. Anal Biochem 1992; 202: 384–9.

33. Colombo G, Clerici M, Garavaglia ME, et al. A step-by-step protocol for assaying protein carbonylation in biological samples. J Chromatogr B Analyt Technol Biomed Life Sci 2016; 1019: 178–90.

34. Calderón-DuPont D, Torre-Villalvazo I, Díaz-Villaseñor A. Is insulin resistance tissue-dependent and substrate-specific? The role of white adipose tissue and skeletal muscle. Biochimie 2023; 204: 48–68.

35. Friedman MI, Sørensen TIA, Taubes G, et al. Trapped fat: Obesity pathogenesis as an intrinsic disorder in metabolic fuel partitioning. Obes Rev 2024; 25: e13795.

36. Hu S, Wang L, Yang D, et al. Dietary Fat, but Not Protein or Carbohydrate, Regulates Energy Intake and Causes Adiposity in Mice. Cell Metab 2018; 28: 415-431.e4.

37. Lackey DE, Lazaro RG, Li P, et al. The role of dietary fat in obesity-induced insulin resistance. American Journal of Physiology-Endocrinology and Metabolism 2016; 311: E989–E997.

38. Barr SB, Wright JC. Postprandial energy expenditure in whole-food and processed-food meals: implications for daily energy expenditure. Food Nutr Res 2010; 54: 5144.

39. Chait A, den Hartigh LJ. Adipose Tissue Distribution, Inflammation and Its Metabolic Consequences, Including Diabetes and Cardiovascular Disease. Front Cardiovasc Med 2020; 7: 22.

40. Baena M, Sangüesa G, Dávalos A, et al. Fructose, but not glucose, impairs insulin signaling in the three major insulin-sensitive tissues. Sci Rep 2016; 6: 26149.

41. Püschel GP, Henkel J. Dietary cholesterol does not break your heart but kills your liver. Porto Biomed J 2018; 3: e12.

42. Chung S, Parks JS. Dietary cholesterol effects on adipose tissue inflammation. Curr Opin Lipidol 2016; 27: 19–25.

43. Caetano AC, da Veiga LF, Capaldi FR, et al. The antioxidant response of the liver of male Swiss mice raised on a AIN 93 or commercial diet. BMC Physiol 2013; 13: 3.

44. Świątkiewicz I, Wróblewski M, Nuszkiewicz J, et al. The Role of Oxidative Stress Enhanced by Adiposity in Cardiometabolic Diseases. Int J Mol Sci 2023; 24: 6382.

45. Arivazhagan L, López-Díez R, Shekhtman A, et al. Glycation and a Spark of ALEs (Advanced Lipoxidation End Products) - Igniting RAGE/Diaphanous-1 and Cardiometabolic Disease. Front Cardiovasc Med 2022; 9: 937071.

46. Corica D, Pepe G, Currò M, et al. Methods to investigate advanced glycation end-product and their application in clinical practice. Methods 2022; 203: 90–102.

47. Magkos F, Fabbrini E, Mohammed BS, et al. Increased whole-body adiposity without a concomitant increase in liver fat is not associated with augmented metabolic dysfunction. Obesity (Silver Spring) 2010; 18: 1510–5.

48. Elsabaawy M. Liver at crossroads: unraveling the links between obesity, chronic liver diseases, and the mysterious obesity paradox. Clin Exp Med 2024; 24: 240.

49. Jakubiak GK, Osadnik K, Lejawa M, et al. Oxidative Stress in Association with Metabolic Health and Obesity in Young Adults. Oxid Med Cell Longev 2021; 2021: 9987352.

50. Blüher M. Metabolically Healthy Obesity. Endocr Rev; 41. Epub ahead of print 1 May 2020. DOI: 10.1210/endrev/bnaa004.

